# Deep learning model fitting for diffusion-relaxometry: a comparative study

**DOI:** 10.1101/2020.10.20.347625

**Authors:** Francesco Grussu, Marco Battiston, Marco Palombo, Torben Schneider, Claudia A. M. Gandini Wheeler-Kingshott, Daniel C. Alexander

**Affiliations:** Queen Square MS Centre, Queen Square Institute of Neurology, Faculty of Brain Sciences, University College London (UCL), UK; Centre for Medical Image Computing, Department of Computer Science, Faculty of Engineering Sciences, University College London (UCL), UK; Philips UK, Guildford, UK; DeepSpin GmbH, Berlin, Germany; Department of Brain and Behavioural Sciences, University of Pavia, Pavia, Italy; Brain MRI 3T Center, IRCCS Mondino Foundation, Pavia, Italy

## Abstract

Quantitative Magnetic Resonance Imaging (qMRI) signal model fitting is traditionally performed via non-linear least square (NLLS) estimation. NLLS is slow and its performance can be affected by the presence of different local minima in the fitting objective function. Recently, machine learning techniques, including deep neural networks (DNNs), have been proposed as robust alternatives to NLLS. Here we present a deep learning implementation of qMRI model fitting, which uses DNNs to perform the inversion of the forward signal model. We compare two DNN training strategies, based on two alternative definitions of the loss function, since at present it is not known which definition leads to the most accurate, precise and robust parameter estimation. In strategy 1 we define the loss as the *l*^2^-norm of tissue parameter prediction errors, while in strategy 2 as the *l*^2^-norm of MRI signal prediction errors. We compare the two approaches on synthetic and 3T *in vivo* saturation inversion recovery (SIR) diffusion-weighted (DW) MRI data, using a model for joint diffusion-T1 mapping. Strategy 1 leads to lower tissue parameter root mean squared errors (RMSEs) when realistic noise distributions are considered (e.g. Rician vs Gaussian). However, strategy 2 offers lower signal reconstruction RMSE, and allows training to be performed on both synthetic and actual *in vivo* MRI measurements. In conclusion, both strategies are valid choices for DNN-based fitting. Strategy 2 is more practical, as it does not require pre-computation of reference tissue parameters, but may lead to worse parameter estimation.

## 1 Introduction

Quantitative Magnetic Resonance Imaging (qMRI) techniques estimate biophysical properties of tissues in each voxel of an MR image [1]. qMRI aims to overcome the limitations of routine clinical imaging, i.e. its lack of sensitivity and specificity to early, widespread pathology that precedes focal lesions. To this end, in qMRI sets of multi-contrast images are acquired by varying the MRI pulse sequence parameters **u** in a controlled fashion. Images are then analysed voxel-by-voxel to estimate **p**, a set of tissue parameters (e.g. diffusion or relaxation properties), via non-linear least square (NLLS) fitting [2]. The estimation effectively inverts the forward model

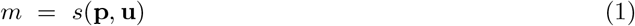

predicting MRI measurements *m* when sequence **u** is used to image tissue with properties **p**. The inversion of the forward model is achieved by minimising an objective function *f*(*m,s*(**p, u**)) with respect to **p**, where *f* quantifies how close signal predictions are to the actual measurements. Maximum-likelihood estimation is achieved if 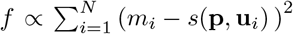 over a set of *N* measurements under the hypothesis of additive Gaussian noise. Alternative definitions of *f* are also in use to account for non-Gaussian noise or in maximum *a posteriori* estimation [3]. Finally, *f* can also include regularisers to stabilise the forward model inversion, as for example terms proportional to **||p||_2_** or ||p||_1_ (Tikhonov or *l^1^* regularisation) [4].

The minimisation of the objective function *f* is typically performed with either Jacobian-based methods, such as Gauss-Newton or Levenberg-Marquardt algorithms, or derivative-free approaches, as for example the Nelder-Mead algorithm [5], which are conveniently available in several computational packages. However, these are inherently slow and often can only find sub-optimal solutions corresponding to local mininima of *f*. Moreover, fitting initialisation also plays a crucial role, and degenerate solutions can be obtained when this is not done accurately [2]. Machine learning has been proposed as an alternative to conventional NLLS to perform qMRI model fitting in qMRI and overcome its limitations. Notable examples include permeability [6] and soma [8] diffusion-weighted (DW) imaging via random forest regression trees, or q-space DW imaging [9] and myelin mapping [4] with deep neural networks (DNNs). Machine learning approaches can be trained off-line and then deployed almost instantaneously when new data come in [4], and offer more stable solutions when trained accurately [6, 7], given their excellent function approximation properties [10].

In this work we introduce an implementation of qMRI fitting based on fully-connected DNNs and use it to compare systematically different training strategies, since at present it is not known which strategy leads to the most accurate, precise and robust parameter estimation. Specifically, we trained DNNs by optimising loss functions defined either i) as the *l*^2^-norm of tissue parameter estimation errors [6, 4, 8], or ii) the *l*^2^-norm of MRI signal estimation errors [7]. We compared the two learning approaches using saturation inversion recovery (SIR) DW data obtained at 3T for joint diffusion and T1 mapping, and performed computational experiments both *in silico* and *in vivo.*

## 2 Methods

In this section we present our implementation of DNN qMRI fitting, which is based on PyTorch and made freely available online (permanent web page: https://github.com/fragrussu/qMRINet). We then introduce two training strategies, and describe experiments performed *in silico* and *in vivo*.

### 2.1 qMRI model fitting with DNNs

Let us consider a qMRI experiment whose aim is the estimation of *P* tissue parameters **p** = [*p*_1_ … *p_P_*]^*T*^ in a voxel according to the generic qMRI model *s*(**p, u**), given a set of *N* ≥ *P* measurements **m** = [*m*_1_ … *m_N_*] performed at sequence parameters **U** = { **u**_1_, …, **u**_*N*_ }. We use a fully-connected DNN to approximate the inverse model **p** ≈ *s*^-1^(**m**, **U**), obtaining an estimator of **p** from the set of input measurements **m**. The DNN input layer is made of *N* neurons, processing the *N*-dimensional array **m**, while the output layer is made of *P* neurons, mapping tissue parameters. Each hidden layer consists of standard linear operators followed by rectified non-linear units (ReLU(*x*) = *max*(0, *x*)). The number of hidden neurons decreases linearly from *N* to *P*, but other architectures are also possible [4, 7]. The DNN is trained with standard back-propagation on synthetic or *in vivo* MRI data. We explore two different training strategies, based on alternative definitions of the training loss function *L* (Figure 1).

**Figure 1:**
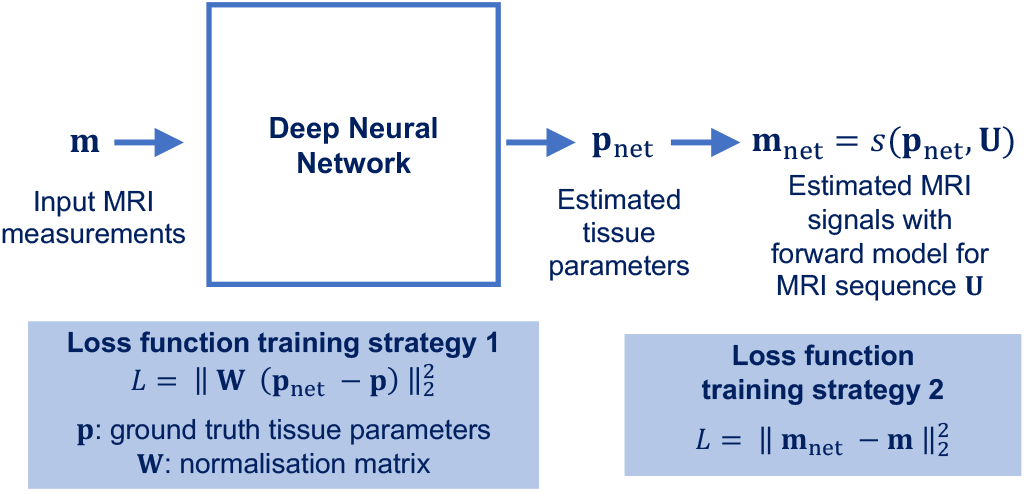
Illustration of our DNN for qMRI model fitting and of its training strategies. Strategy 1: loss defined on tissue parameters. Strategy 2: loss defined on MRI signals.

#### 2.1.1 Training strategy 1: loss defined on tissue parameters

The loss *L* is defined as the squared *l*^2^-norm of tissue parameter estimation errors:

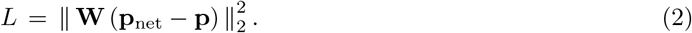

Above, **p**_net_ ≈ *s*^−1^(**m, U**) is the DNN output, **p** are ground truth tissue parameters and 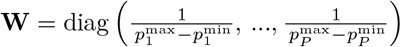 is a normalisation matrix controlling for the fact that parameters can be defined over different numerical ranges (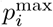 and 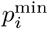 are the upper and lower bound of the *i*-th parameter for *i* = 1, …, *P*). Examples of paired measurements/parameters training examples can be synthesised with model *s*(**p, u**), or gathered from previous NLLS fitting.

#### 2.1.2 Training strategy 2: loss defined on signals

The loss *L* is defined as the squared *l*^2^-norm of signal prediction errors, similarly to routine NLLS methods:

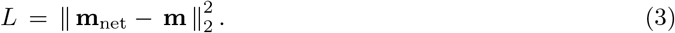

In Equation 3, **m** is the *N*-dimensional array of input qMRI measurements, while **m**_net_ is an estimate of **m** obtained by re-applying the forward model *s* to the DNN output **p**_net_ ≈ *s*^-1^(**m, U**), i.e.

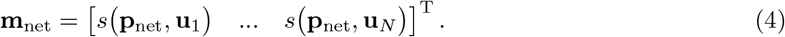

In practice, an additional normalisation is performed to map the the *i*-th output neuron activation *v_i_* to tissue parameter *p_i_* before computing Equation 4, i.e.

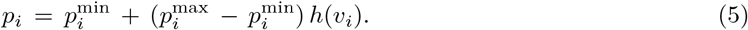

Above, 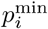 and 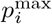 are again the lower/upper bounds of *p_i_*, while *h*(*v_i_*) is

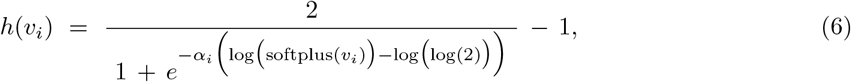

where softplus(*x*) = log(1 + *e^x^*) and where *α_i_* is an actual network parameter that is learnt during training, similarly to all linear layer weights and biases. The mapping in Equation 5 prevents infinite or not-a-number values to occur when computing Equation 4. Training is performed on synthetic or actual qMRI measurements and no knowledge of ground truth tissue parameters is required.

### 2.2 *In silico* study

#### 2.2.1 Signal synthesis

We performed numerical simulations to compare DNN fitting implemented according to strategies 1 and 2. To this end, we consider one among several potential qMRI signal models and focus on increasingly popular diffusion-relaxation imaging (DRI), which exploits the complementary information from different qMRI contrasts in a unified acquisition [11, 12]. Specifically, we consider the case of joint diffusion and T1 mapping in the brain, as achievable in inversion recovery (IR) [13] or saturation IR (SIR) [14] DWI. We borrow a previously proposed tensor approach [13] and adapt it to directionally-averaged (i.e. spherical mean) DW signals, as this removes the dependence on the underlying fibre orientation distribution [15]. The forward model *s*(**p, u**) is

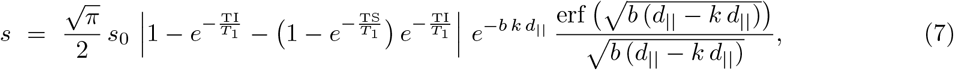

with tissue parameters **p** = [*d*_||_ *k T s*_0_]^*T*^ (parallel diffusivity *d*_||_, defined in [0.01; 3.2] *μ*m^2^ms^−1^; anisotropy parameter *k*, defined in [0; 1] and such that the perpendicular diffusivity *d*_⊥_ is *d*_⊥_ = *k d*_||_(anisotropy *k* ∝ 1/*k*); longitudinal relaxation time *T*_1_, defined [100; 4000] ms; apparent proton density *s*_0_, defined in [0.5; 5]) and sequence parameters **u** = [TS TI *b*] (saturation-inversion delay TS; inversion-excitation delay, i.e. inversion time TI; b-value *b*). Tissue parameters are defined within biologically plausible ranges at 3T.

We generated synthetic SIR DWI signals with Equation 7 (160,000 as training set; 40,000 as validation set; 16,000 as test set). For this, we used tissue parameters drawn from uniform distributions, using the same parameter bounds reported above. A protocol made of *N* = 32 measurements was used, replicating acquisitions performed *in vivo* and described in sub-section 2.3. These were: (*b,* TI) = {0, 1000, 2000, 3000}smm^−2^ × {70, 320, 570, 820, 1070, 1320, 1570, 1820} ms, with TS fixed to TS = 300 ms. Signals were corrupted with both Gaussian and Rician noise, varying the signal-to-noise ratio (SNR) across voxels uniformly within the range 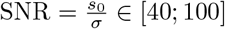.

#### 2.2.2 DNN fitting

qMRI measurements **m** = [*m*_1_(*b*_1_, TI_1_) … *m*_32_(*b*_32_, TI_32_)]^T^ from each synthetic voxel were normalised as 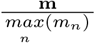, and then used to train DNNs according to both strategy 1 and 2. DNNs consisted of 7 hidden layers with input/output sizes of (32 × 28), (28 × 24), (24 × 20), (20 × 16), (16 × 12), (12 × 8), (8 × 4), and were trained with ADAM [16] for 250 epochs (learning rate: 0.0005) grouping voxels in mini-batches (one DNN update per mini-batch of 100 voxels). We repeated training 8 times, each time with a different random DNN initialisation, to minimise the risk of incurring in local minimina. The DNN providing the lowest validation loss was used to predict tissue parameters and signals on the test data. Root mean squared error (RMSE) with respect to synthetic noisy signals and ground truth tissue parameters were computed.

### 2.3 *In vivo* study

#### 2.3.1 MRI acquisition

We performed brain DRI on a 3T Philips Ingenia CX system. Scans were performed after obtaining informed written consent, and were approved by a local Research Ethics Board. We recruited 3 healthy volunteers (2 females) and used a multi-slice SIR [14] DW EPI sequence, with a 32-channel head coil for signal detection. Sequence parameters were: 48 axial slices, 2.4mm-thick; field-of-view: 230 × 230mm^2^; in plane resolution: 2.4 × 2.4mm^2^; TR = 2563 ms; TE = 90 ms; TS = 300 ms; SENSE = 2; MB = 3; 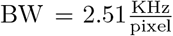. 528 images were acquired with 32 unique (*b*, TI) values ((*b,* TI) = {0, 1000, 2000, 3000} smm^−2^ × {70, 320, 570, 820, 1070, 1320, 1570, 1820} ms, 21 directions for non-zero *b*, 3 images for each *b* = 0) in 46 minutes, which included an image with reversed phase encoding for EPI distortion mitigation.

#### 2.3.2 MRI post-processing

We denoised scans with the MP-PCA technique [17], subsequently mitigating noise floor [18] and Gibbs ringing [19]. We corrected for motion and eddy current via affine co-registration based on NiftyReg [20] and EPI distortions with FSL [21], and obtained a brain mask [22]. Finally, we averaged images corresponding to different gradient directions at fixed (*b,* TI), obtaining a data set made of *N* = 32 unique (*b*, TI) measurements.

#### 2.3.3 DNN fitting

The DNNs trained for *in silico* experiments with Rician noise were also used for fitting Equation 7 on *in vivo* data. Moreover, for training strategy 2, we also trained a DNN on actual *in vivo* measurements extracted from within the brain mask (which included cerebrospinal fluid (CSF)), using the same learning settings described in section 2.2.2. We followed a leave-one-out strategy, training on 2 subjects and using the DNN for model fitting on the other (number of training voxels comparable to *in silico* training). Fitting provided voxel-wise tissue parameter maps and MRI signal estimates, from which RMSEs with respect to actual *in vivo* MRI measurements were computed.

## 3 Results

Figure 2 shows predictions of MRI signals and of tissue parameters for simulations conducted with Gaussian noise. Inspection of the plots suggests that strategy 1 and 2 perform similarly. However, when Rician noise is considered (Figure 3), strategy 1 enables more accurate tissue parameter estimation. Nonetheless, strategy 2 still provides a better estimation of the MRI signal as compared to strategy 1, which underestimates slightly the signal level.

**Figure 2:**
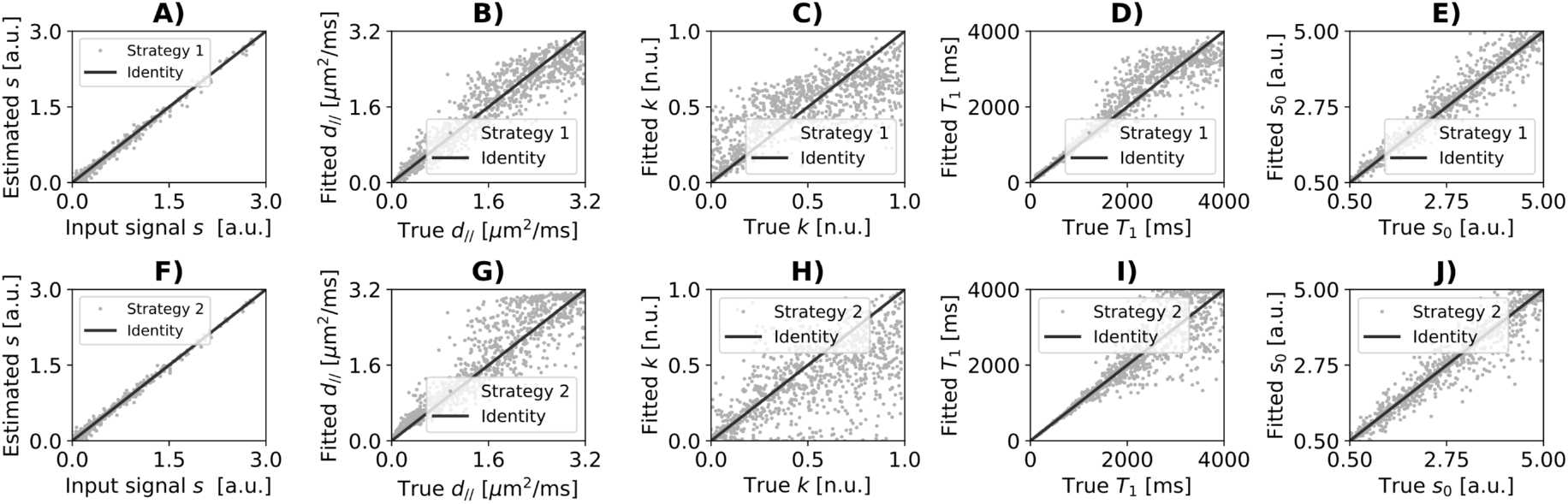
Scatter plots of simulation results obtained on the test set when Gaussian noise is considered. Panels A and F show predicted signal (y-axis) against input measurements (x-axis). Panels B-E and G-J show fitted parameters (y-axis) against ground truth tissue parameters (x-axis) (*d*_∥_ in B, G; *k* in C, H; *T*_1_ in D, I; *s*_0_ in E, J). Top (A-E): training strategy 1. Bottom (F-J): training strategy 2.

**Figure 3:**
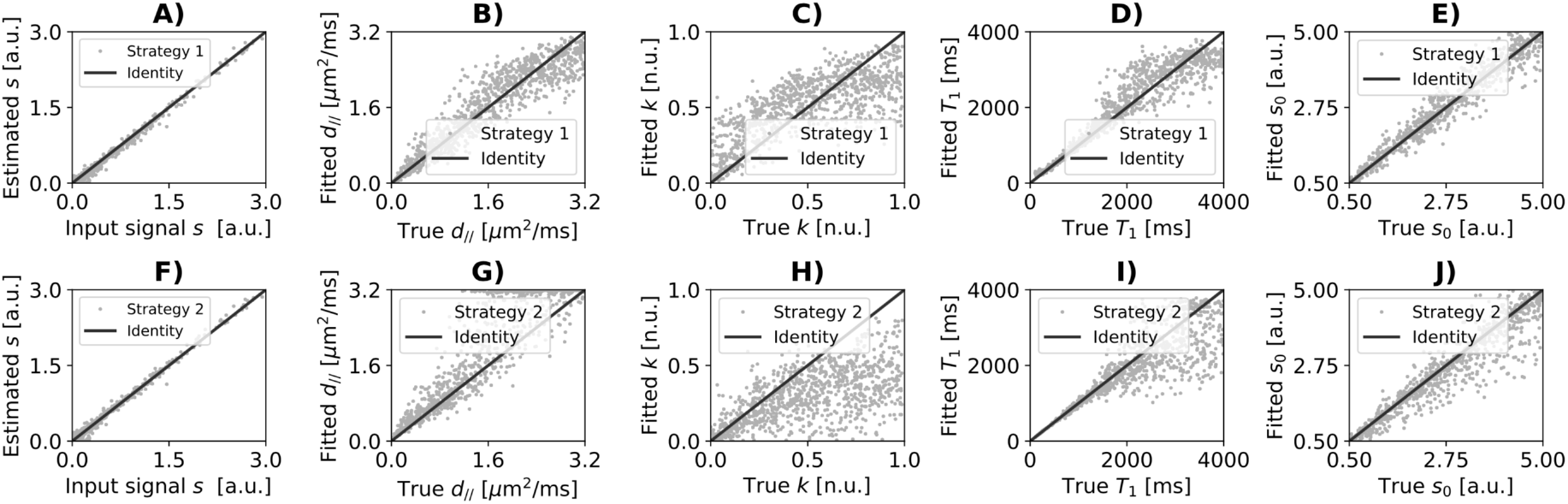
Scatter plots of simulation results obtained on the test set when Rician noise is considered. As in Figure 2, panels A and F show predicted signal against input measurements, while panels B-E and G-J fitted against ground truth tissue parameters.

Table 1 reports statistics of signal and tissue parameter RMSE distributions obtained on the test set with both Gaussian and Rician noise. Strategy 2 provides lower median RMSE for almost all tissue parameters as well as for the signal estimation than strategy 1 when Gaussian noise is considered. However, it also provides wider RMSE distributions. With Rician noise instead, strategy 1 provides overall lower RMSE figures for tissue parameter estimates, while strategy 2 still provides lower RMSE for MRI signal predictions. For both strategies, Rician noise leads to worse performances compared to Gaussian noise.

**Table 1:**
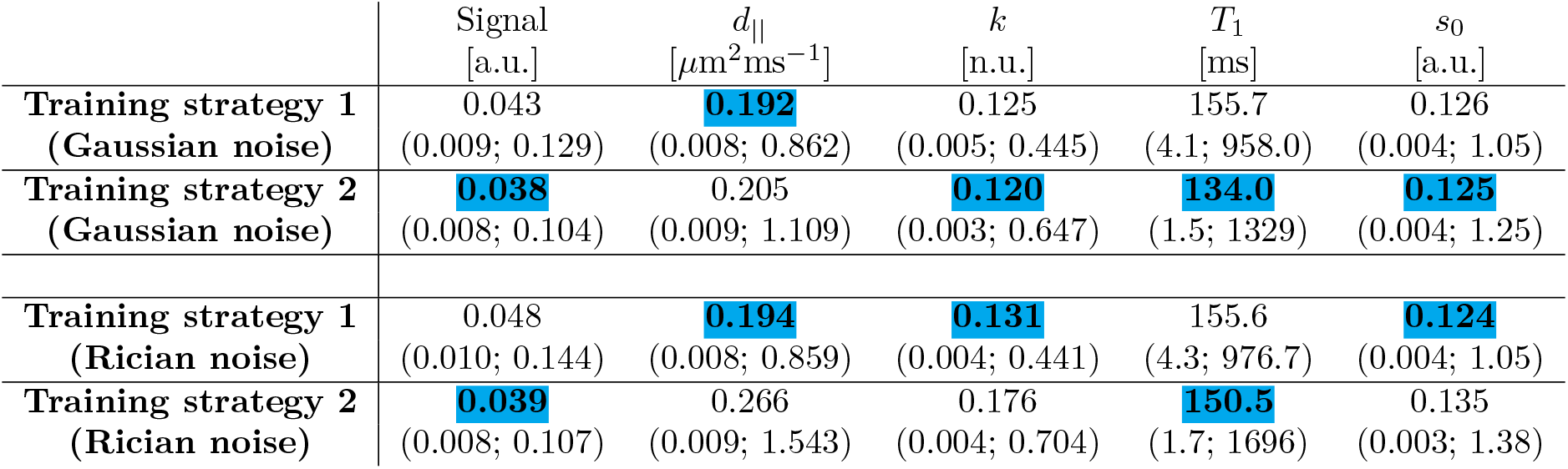
Root mean squared errors (RMSEs) between MRI measurements and signal predictions as well as between ground truth and fitted parameters from simulations. Median and 2.5-97.5 percentiles (in brackets) over the test set are reported (lowest median RMSE highlighted in light blue). Results from simulations conducted with both Gaussian and Rician noise cases are reported.

Figure 4 shows examples of voxel-wise tissue parameter maps obtained *in vivo.* The two training strategies provide metrics that are qualitatively similar and that exhibit similar between-tissue contrasts. However, some differences between the approaches are seen (e.g. slightly higher *d*_∥_ and lower *k* in strategy 2 compared to strategy 1), especially in areas with strong CSF partial volume. For training strategy 2, training the DNN on synthetic signals or actual *in vivo* measurements essentially provides the same results.

**Figure 4:**
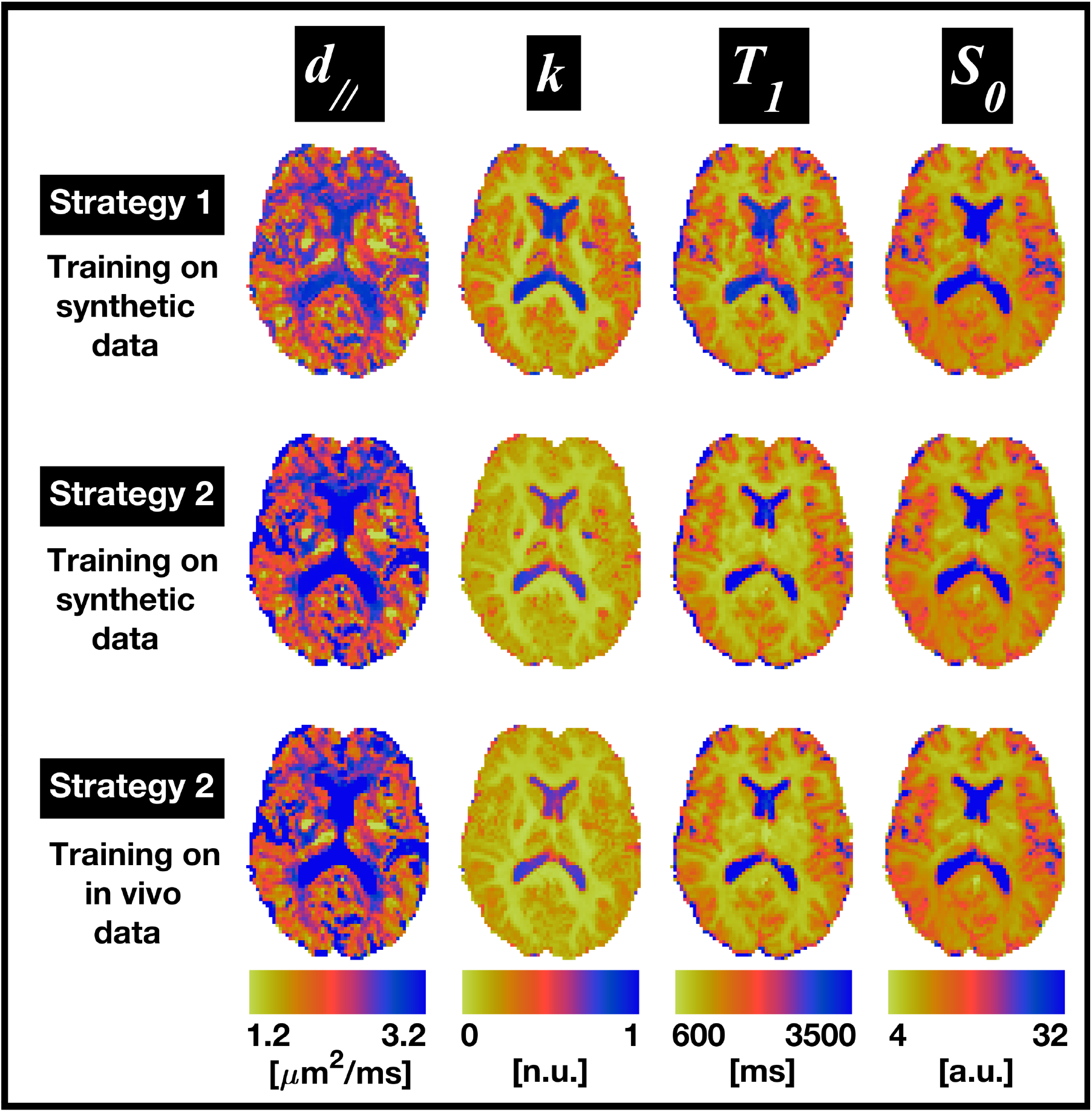
Examples of *in vivo* parametric maps. From left to right: *d*_∥_, *k*, *T*_1_ and *s*_0_. Top row: strategy 1. Central row: strategy 2, with DNN trained on synthetic MRI signals. Bottom row: strategy 2, with DNN trained on actual *in vivo* MRI measurements.

Figure 5 shows examples of images acquired *in vivo* as well as DNN predictions (leave-one-out experiment). All training strategies enable learning of a variety of diffusion and T1 contrasts, as predictions are qualitatively similar to the acquired images. On visual inspection, predictions from strategy 2 more closely resemble *in vivo* images as compared to strategy 1.

**Figure 5:**
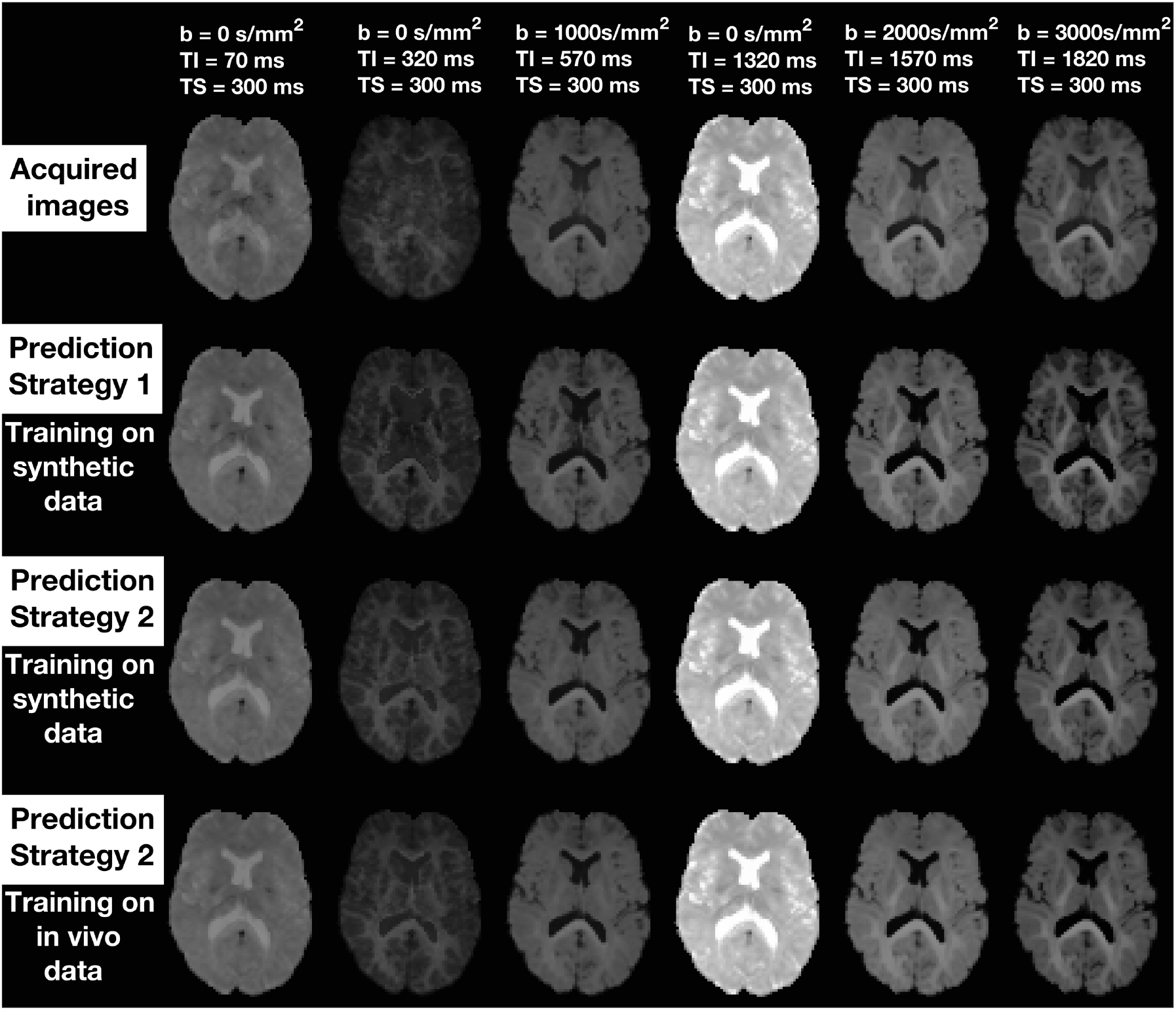
Examples of images acquired *in vivo* and DNN predictions. Left to right: different contrasts (spherical mean images; sequence parameters reported on top). Top to bottom: acquired images; images predicted by DNN trained with strategy 1; images predicted by DNN trained with strategy 2 on synthetic signals; images predicted by DNN trained with strategy 2 on *in vivo* signals. The same grey scale color scheme has been used for all images (intensities are comparable across contrasts/strategies).

Finally, Table 2 reports signal prediction RMSE as measured on *in vivo* data. Results are in line with RMSE figures from simulations. In all cases, training strategy 2 has slightly lower RMSE compared to training strategy 1. For training strategy 2, the lowest RMSE figures are obtained when training on synthetic data. However, training on actual *in vivo* measurements yields almost identical performances (difference of median RMSE less than 1%).

**Table 2:** *In vivo* root mean squared error (RMSEs) between MRI measurements and signal predictions. Median and 2.5-97.5 percentiles (in brackets) obtained within the brain are reported, highlighting in light blue the approach providing the lowest median RMSE.

| MRI signal<br>RMSE [a.u.] | Strategy 1<br>(train on syn. data) | Strategy 2<br>(train on syn. data) | Strategy 2<br>(train on *in vivo* data) |
| --- | --- | --- | --- |
| Volunteer 1 | 0.208<br>(0.078; 0.674) | **0.142**<br>(0.061; 0.626) | 0.143<br>(0.060; 0.640) |
| Volunteer 2 | 0.191<br>(0.070; 0.589) | **0.125**<br>(0.056; 0.519) | 0.126<br>(0.054; 0.524) |
| Volunteer 3 | 0.233<br>(0.086; 0.742) | **0.164**<br>(0.068; 0.665) | 0.165<br>(0.067; 0.675) |

## 4 Discussion

In this work we studied DNN-based qMRI signal model fitting. We used DNNs to implement an estimator of tissue parameters from input MRI measurements, and ran experiments on synthetic and *in vivo* DRI data. Specifically, we considered a model describing directionally-averaged SIR DWI data for joint diffusion and T1 mapping. We performed computational experiments on synthetic and *in vivo* scans, which were acquired at 3T on 3 healthy volunteers. Both *in silico* and *in vivo* data were used to explore two alternative training strategies: in strategy 1, the training loss is defined as the *l^2^*-norm of tissue parameter estimation errors; in strategy 2, as the *l*^2^-norm of MRI signal reconstruction errors.

Our results suggest that both strategies are viable options for DNN-based fitting. When Gaussian noise is considered, the performances of the two strategies are comparable: strategy 2 provides higher accuracy than strategy 1 in parameter estimation, at the expenses of lower precision. However, when more realistic noise distributions (i.e. Rician) are considered, the accuracy of strategy 1 is superior to strategy 2. We speculate that such reduction in accuracy for strategy 2 results from the fact that DNNs trained end up learning features that originate from the noise floor, adjust tissue parameter accordingly to accommodate for this. Moreover, noise floor bias could also explain the fact that the MRI signal is slightly underestimated in strategy 1 when Rician noise is considered.

Importantly, it should be noted that unlike in simulations, one does not have access to ground truth tissue parameters *in vivo.* Therefore, care is needed when extrapolating the better parameter estimation obtained in strategy 1 from simulations to the *in vivo* case.

Another key observation relates to the fact that strategy 1 requires paired examples of MRI measurements and corresponding tissue parameters. Such examples could be gathered from previous NLLS fitting, or synthesised *in silico*, as done here. The former is anything but convenient, while the latter would lead to assumptions on the underlying noise level and distribution. Such assumptions can instead be avoided in strategy 2, as demonstrated here when training is performed on actual *in vivo* measurements, obtaining robust fitting results.

Finally, we acknowledge a number of potential limitations. Firstly, we considered only one qMRI signal model. In future, we will assess the generalisability of our findings by considering additional models. Secondly, we did not compare our fitting to non-DNN-based alternatives. While these are certainly interesting to put DNN performance in context, they are beyond the scope of this work, since here we focus on specific design choices of DNN-based fitting. Thirdly, we acknowledge that residual noise floor bias may still affect *in vivo* MRI measurements even after Rician bias mitigation. In future, we will explore more effective noise floor correction techniques. Furthermore, in our comparative study some tissue parameters (e.g. *s*_0_) were estimated better than others (e.g. *k*). This may be a result of the intrinsic poor sensitivity of the MRI protocol used here with respect to certain parameters. In future we will overcome this limitation by optimising the (*b,* TI) sampling scheme. Finally, we acknowledge that it cannot be concluded whether our results hold in pathology, e.g. in small focal lesions affecting only a small percentage of the voxels in the image, since here we only consider healthy subjects. We speculate that in those cases strategy 1 may provide additional advantages as compared to strategy 2 in the under-represented lesion, especially if training is performed on actual *in vivo* measurements. We will address this question in future work.

## 5 Conclusion

DNN-based fitting is a viable approach for brain DRI modelling. Training by minimising parameter estimation errors or signal reconstruction errors are both valid choices. The latter is more practical as it does not require pre-calculation or synthesis of tissue parameters, but may provide worse parameter estimation.

## 6 Acknowledgements

This project was funded by the Engineering and Physical Sciences Research Council (EPSRC EP/R006032/1, M020533/1, EP/N018702/1 G007748, I027084) and by the UK Research and Innovation (MR/T020296/1, funding M.P.). This project has received funding under the European Union’s Horizon 2020 research and innovation programme under grant agreement No. 634541, and from: Rosetrees Trust (UK, funding F.G.); Spinal Research (UK), Wings for Life (Austria), Craig H. Neilsen Foundation (USA) for INSPIRED; Wings for Life (169111); UK Multiple Sclerosis Society (grants 892/08 and 77/2017); the Department of Health’s National Institute for Health Research (NIHR) Biomedical Research Centres and UCLH NIHR Biomedical Research Centre.

## 7 Declaration of conflict or commercial interest

T.S. worked for Philips (UK) and now works for DeepSpin (Germany).

## Notes

https://github.com/fragrussu/qMRINet

### Competing Interest Statement

Torben Schneider worked for Philips (UK) and now works for DeepSpin (Germany).

